# Helical remodeling augments 5-lipoxygenase activity

**DOI:** 10.1101/2022.03.31.486615

**Authors:** Eden M. Gallegos, Tanner D. Reed, Forge A. Mathes, Nelson Guevera, David B. Neau, Wei Huang, Marcia E. Newcomer, Nathaniel C. Gilbert

## Abstract

The synthesis of pro-inflammatory leukotrienes implicated in asthma, allergic rhinitis, and atherosclerosis is initiated by the enzyme 5-lipoxygenase (5-LOX). The crystal structure of human Stable-5-LOX revealed a conformation where the catalytic iron was inaccessible to bulk solvent as two aromatic residues on a conserved helix-α2 (Hα2) plugged the substrate access portal. Here, we present a new conformation of 5-LOX where Hα2 adopts an elongated conformation equivalent to that described in other animal lipoxygenase structures. The sigmoidal kinetic behavior of 5-LOX, which is indicative of positive cooperativity, is consistent with a substrate-induced conformational change that shifts the ensemble of enzyme populations to favor the catalytically competent state. Strategic point mutations along Hα2 designed to unlock the closed conformation and elongate Hα2 resulted in improved kinetic parameters, altered limited-proteolysis data, and a drastic reduction in the length of the lag phase yielding the most active 5-LOX enzyme to date. Structural predictions by AlphaFold2 of these variants statistically favor an elongated Hα2 and reinforce a model in which improved kinetic parameters correlate with a more readily adopted, open conformation.

## Introduction

The enzyme 5-lipoxygense (5-LOX) performs the initial step in the synthesis of leukotrienes (LTs), potent inflammatory mediators linked to asthma, allergic disorders, and atherosclerosis^1-3^. The substrate arachidonic acid (AA) is derived from the nuclear membrane and presented to 5-LOX by its partner protein 5-lipoxygenase-activating protein (FLAP)^4,5^. The production of LTs begins when cells are stimulated to release Ca^2+^, triggering the translocation of cytosolic phospholipase A_2_ and 5-LOX to the nuclear membrane^6,7^. Phospholipase A2 cleaves AA from membrane phospholipids and FLAP-bound AA is accessed by 5-LOX, which then transforms it to leukotriene A_4_ (LTA_4_) in a two-step reaction (for review see^8^). 5-LOX is unique to the lipoxygenase family in that it is the only isoform known to require a helper protein^4^ (FLAP) and perform a second reaction, the leukotriene A_4_ synthase activity^9^. The integral membrane protein FLAP is required for the efficient generation of the LT product in intact cells; as such, many anti-LT therapies in development focus on antagonists to FLAP (for review see^10^). A high dependence on FLAP for effective LTA_4_ production suggests there are interactions between 5-LOX and its partner that are essential for the enzyme to achieve full activity^11^.

Traditional approaches to understanding enzyme-substrate interactions were first described by the inflexible “lock and key” model of Emil Fischer in 1894, which further evolved to an “induced fit” model by Koshland^12^. Still, there remains much to understand about the types of conformational changes enzymes may undergo before they achieve full catalytic proficiency (for review see^13^). Regulation as a consequence of conformational changes transduced to the active site are the basis for both allostery and cooperativity^14^. Cooperativity is easily envisioned for oligomeric enzymes given an obvious communication route between distinct active sites, but cooperativity has been observed for monomeric enzymes with single substrate binding sites (for review^15^). In such a case with positive cooperativity, the presence of increasing amounts of substrate invokes a shift of the population of ensembles towards a fully activated-enzyme conformation. A sigmoidal response to increasing substrate concentration yields an ultrasensitive, “switch-like” response over a narrow range of substrate concentrations^16^. One can envision that the regulation in the production of the highly potent LTs would benefit from such a mechanism. In an effort to understand how substrate might enter the closed active site of 5-lipoxygenase, we pursued crystal structures of the enzyme in the presence of substrate and inhibitors and identified an “open” conformation of the enzyme. Kinetic data presented here support a model of cooperative regulation of the monomeric enzyme where the presence of substrate can shift the balance in the conformational ensemble of the monomeric enzyme towards a fully activated enzyme.

5-LOX is one of several lipoxygenase (LOX) isoforms expressed in animals^17^. The enzymes share a common structural framework and a highly conserved catalytic core yet differ in product specificity (for review^18^). However, one striking structural difference of 5-LOX is an alternate conformation of an α-helix that rims the active site: helix-α2 (Hα2)^19^. An elongated Hα2 with at least 6 turns is observed in the structures of coral 8*R*-lipoxygenase, human 15-lipoxygenase-2, porcine 12-lipoxygenase, and rabbit 15-lipoxygenase, while the counterpart in 5-LOX is a broken helix with segments of at most three-helical turns that form a v-like structure (Fig. 1). While the details of a putative interaction between 5-LOX and the membrane or FLAP are lacking, aromatic side chains from Hα2 as positioned in 5-LOX obstruct the passage of AA into the active site. Thus, the “closed” Hα2 structure of 5-LOX may play a key regulatory role in the enzyme’s activity by limiting access to the catalytic iron. Moreover, in this closed state the side chains of multiple aromatic amino acids (F177, Y181, F193, F197) distributed along Hα2 are buried. Were 5-LOX able to adopt the commonly observed configuration of Hα2, these aromatic amino acids would become exposed and potentially require burial in a hydrophobic environment^20^.

**Fig. 1.**
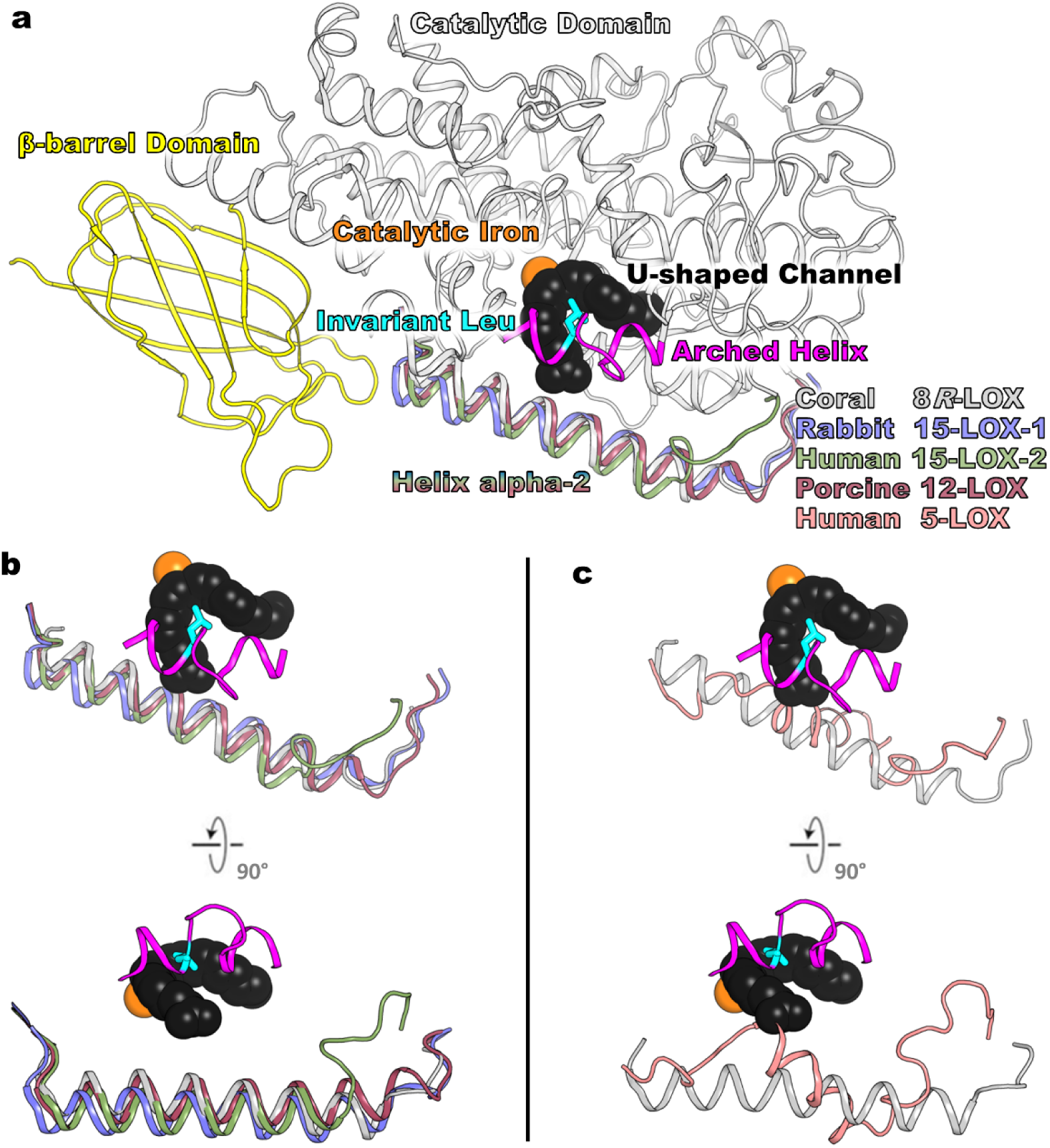
Common architecture of animal lipoxygenases. **a**, Animal lipoxygenases are two domain structures: N-terminal β-barrel domain in yellow and mostly α-helical catalytic domain in grey ribbon. Active site iron is shown as orange sphere. The magenta arched helix with the invariant Leu (c, cyan) spans over the U-shaped channel (black space filling from structure of AA bound to 8R-LOX). **b**, The Hα2s from Coral 8*R*-LOX (grey, 4QWT.pdb), Rabbit 15-LOX-1 (blue, 2P0M.pdb), Human 15-LOX-2 (green, 4NRE.pdb), and Porcine 12-LOX (raspberry, 3RDE.pdb) are one long helix that rims the active site. **c**, Human 5-LOX (salmon, 3O8Y.pdb) Hα2 zig-zags along the body of the protein and blocks entrance to the U-shaped channel.

We report here an elongated Hα2 in crystal structures of 5-LOX determined with data from crystals soaked with common 5-LOX inhibitors or substrate. We asked whether such a conformational change might control access to the active site and play a role in the regulation of 5-LOX activity. To test our model, we engineered point mutations along Hα2 to favor the elongated, open conformation observed in other animal LOXs based in part on our crystal structures of an open conformation of 5-LOX. In this report, we show that key residues that “plugged” the active site in earlier structures are displaced ∼6 Å from their positions in the broken-helix conformation. Kinetic, computational, and limited proteolysis data with variants designed to favor the elongated helix and an open conformation support our idea that the commonly observed open LOX conformation is available to 5-LOX, as well. Further, we show that opening access to the catalytic site leads to increased enzymatic activity without impacting product fidelity. Lastly, we performed a post-hoc analysis and found that the new protein-folding program AlphaFold2 (AF2) predicts “opened” structures for these variants; to our knowledge, this is the first example of point mutations significantly altering the structure prediction by AF2. We propose that this “open” model of 5-LOX may have structural elements in common with the membrane-bound activated enzyme interacting with its partner FLAP.

## Results

### Crystal structures exhibit an elongated Hα2 for Stable-5-LOX

The crystal structure of Stable-5-LOX revealed an overall architecture similar to other animal LOXs: an amino terminal β-barrel domain (residues 1-112) that confers membrane binding, and the larger, primarily α-helical domain (residues 113-673) that harbors the catalytic iron. In most animal LOX structures, a U-shaped cavity lined with hydrophobic amino acids that is complementary in shape to the four *cis* double bonds of the substrate is clearly visible (Fig. 1)^18^. Another prominent feature of LOX structure is the “arched helix” that shelters the U-shaped cavity and contributes an invariant Leu (5-LOX, 414) that pinpoints the key pentadiene on AA for hydrogen abstraction. A striking structural difference observed between Stable-5-LOX and other animal LOXs, some co-crystallized with either substrate or a competitive inhibitor, lies in Hα2 of the catalytic domain. The Stable-5-LOX structure was revealed in a “closed” conformation with the catalytic iron inaccessible to bulk solvent and the U-shaped cavity plugged by F177 and Y181 (FY plug), which protrude from this broken Hα2 (Fig. 1).

We sought to determine structures that might give insight into how the catalytic iron is accessed by soaking Stable-5-LOX crystals with various inhibitors or substrate. Electron density maps calculated with data from 11 distinct soaking experiments (of >30 total datasets collected) revealed density for an elongated Hα2 (Fig. 2b) despite an absence of ligand or substrate density. The “broken” Hα2 in these structures is propagated in a contiguous fashion in one of the protomers to include amino acids 171-197 (Fig 2b). Simultaneously, Hα2 of the other protomer in the asymmetric unit becomes highly disordered, with even the main chain carbons untraceable. Weak electron density is also observed for the arched helix of both protomers with low real-space correlation coefficients (RSCC), so these peptide regions are left unmodelled in the final structures (see methods sections for details on omitting peptides). We focus here on two representative data sets that provide structures at 2.1 Å (PDB code: 7TTJ, two protomers in the asymmetric unit) and 2.43 Å resolution (PDB code: 7TTL, four protomers in the asymmetric unit) with a remodeled Hα2. Data collection and refinement statistics can be found in Supplementary Table 1.

**Fig. 2.**
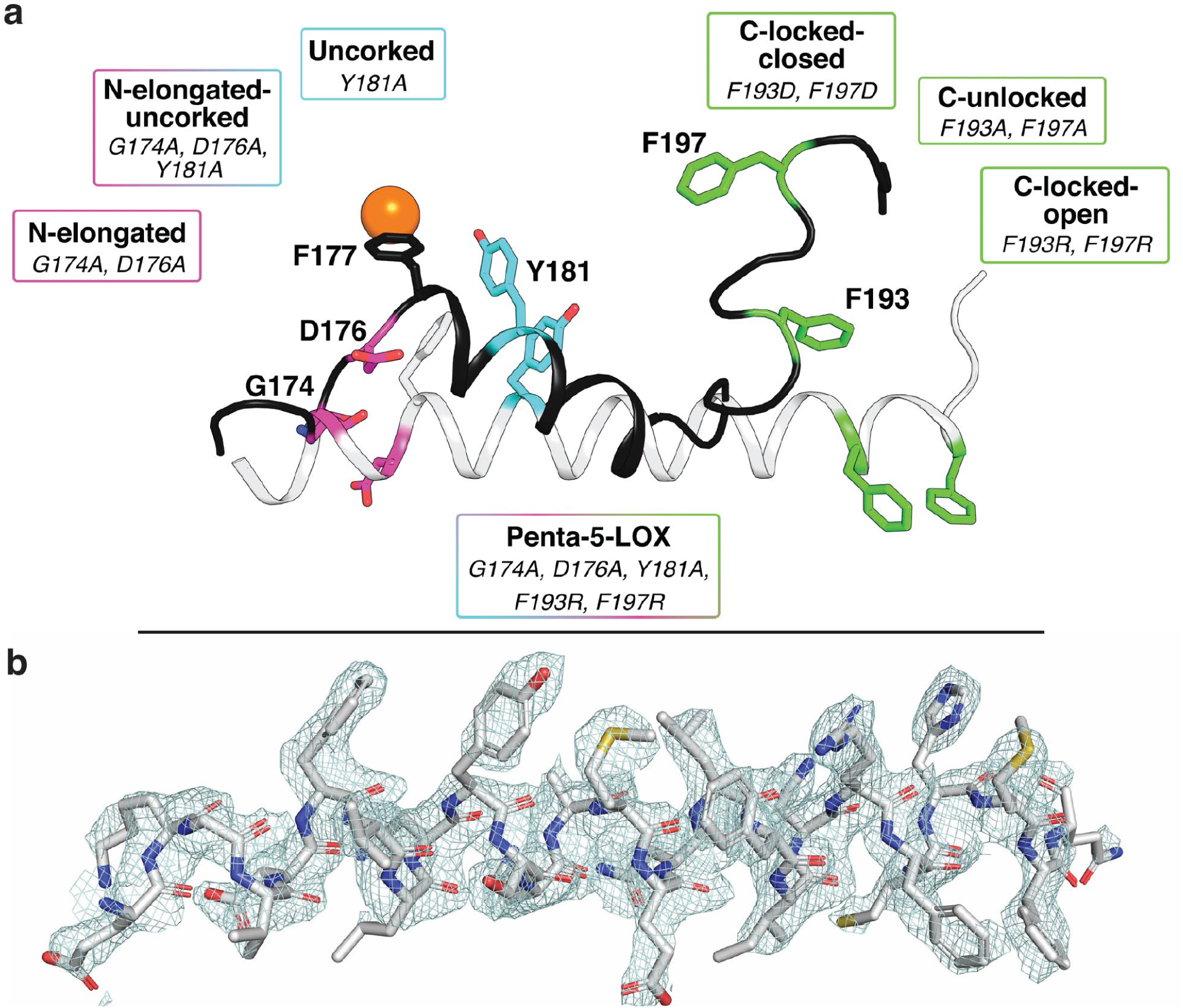
Schematic overview of active and resting models of Stable-5-LOX created through mutagenesis and electron density of an elongated Hα2. **a**, Hα2 of 5o8y.pdb is depicted in black as a cartoon with residues of interest shown as sticks. The alternate conformation of Hα2 is colored white. Pink colored sticks indicate amino-terminal mutated residues, light blue indicates the corking residue that was mutated, and green indicates carboxy-terminal residues that were mutated. F177 is shown as a stick. The active site iron is colored orange. **b**, Electron density |2Fo –Fc| contoured to 1σ of the elongated version of Hα2 with residues depicted as sticks. Atoms are colored as follows: grey, carbon; red, oxygen; blue, nitrogen; yellow, sulfur.

Elongated Hα2 adopts a conformation similar to other “open” LOX structures and extends along the surface of the catalytic domain (Fig. 2). Residues F177 and Y181 that plugged the active site in the closed conformation undergo a main chain shift of 7.2 Å and 4.2 Å, respectively (Supplementary Fig. 2). A coordinated rotamer switch of these aromatic residues positions the side chains away from the body of the enzyme while maintaining the pi-pi stacking observed in the closed conformation (all interactions were defined by the RING 2.0 web server^21^). Another notable displacement is that of D176, which participates in an intricate network of side-chain to main-chain and side-chain to side-chain H-bonds as well as ionic interactions with L179, Q413, and K409 when the substrate portal is closed. The main chain of D176 undergoes an 8.2 Å displacement between these two states. A loop-to-helix transition elongates the C-terminal end of Hα2 an additional 2 turns, with the α-carbons of F193 and F197 repositioning at distances of 9.7 Å and 22.6 Å, respectively (Supplementary Fig. 2). These Phe residues are buried in the body of the enzyme when in a closed conformer, but in the open structure protrude from the catalytic body to bury into the neighboring protomer of the asymmetric unit.

### Molecular dynamic ensemble refinement of Stable-5-LOX

Our repertoire of structural studies has clearly revealed backbone flexibility, and even remodeling, in the segment of the enzyme that includes Hα2. Hα2 has been observed fully disordered^22^, broken and segmented^19^, partially disordered^23^, and now elongated and continuous. We asked what features of the structure are key to a transition from a closed structure to a conformation capable of binding substrate^24^. Molecular Dynamic (MD) simulations that sample both local atomic fluctuations and rigid-body movement by a <u>T</u>ranslation-<u>L</u>ibration-<u>S</u>crew-rotation (TLS) model all time-averaged restrained to the X-ray data were pursued. We performed an ensemble refinement on the two highest resolution structures collected to date of both the closed (PDB code: 7TTK) and open (PDB code: 7TTJ) conformations at 1.98 Å and 2.10 Å, respectively. Ensemble refinement can help illustrate the conformational heterogeneity of protein structure but cannot help model highly disordered regions with low RSCC. As expected, the β-barrel domain was calculated to have a high divergence of backbone positions in the ensemble models, which is especially notable in the membrane-binding loop regions. A similar observation was first detailed in the ensemble structure of bovine pancreatic phospholipase A2 in the loops of the β-barrel domain where it peripherally associates with the lipid bilayer^25^. In the catalytic domain, the main-chain atoms of the N-terminal portion of Hα2 had strikingly dissimilar ensembles in both the closed and open structures (Fig. 3). In contrast, the ensembles of the plugging residues F177/Y181 in the closed structure superimpose exceedingly well and report a low B-factor, likely indicative of a preferred conformation in aqueous buffer. For comparison, this same plug in the open structure has higher computed B-factors with the F177 sampling many rotomeric positions and backbone movement. This Phe is highly conserved in animal LOXs, and its displacement is requisite for activity. At the C-terminus of Hα2, the aromatic residues F193/F197 sample fewer conformations in the ensemble models as the FY plug. The overall pattern for high B-factors is mostly restricted to one face of the protein where the membrane-binding loops of the β-barrel domain and Hα2 of the catalytic domain lie. Interestingly, this face of 5-LOX is predicted to interact with the membrane by the Position of Protein in Membranes (PPM) server^20^, a prediction consistent with the dynamic nature of a peripheral membrane protein that undergoes a phase transition^26^.

**Fig. 3.**
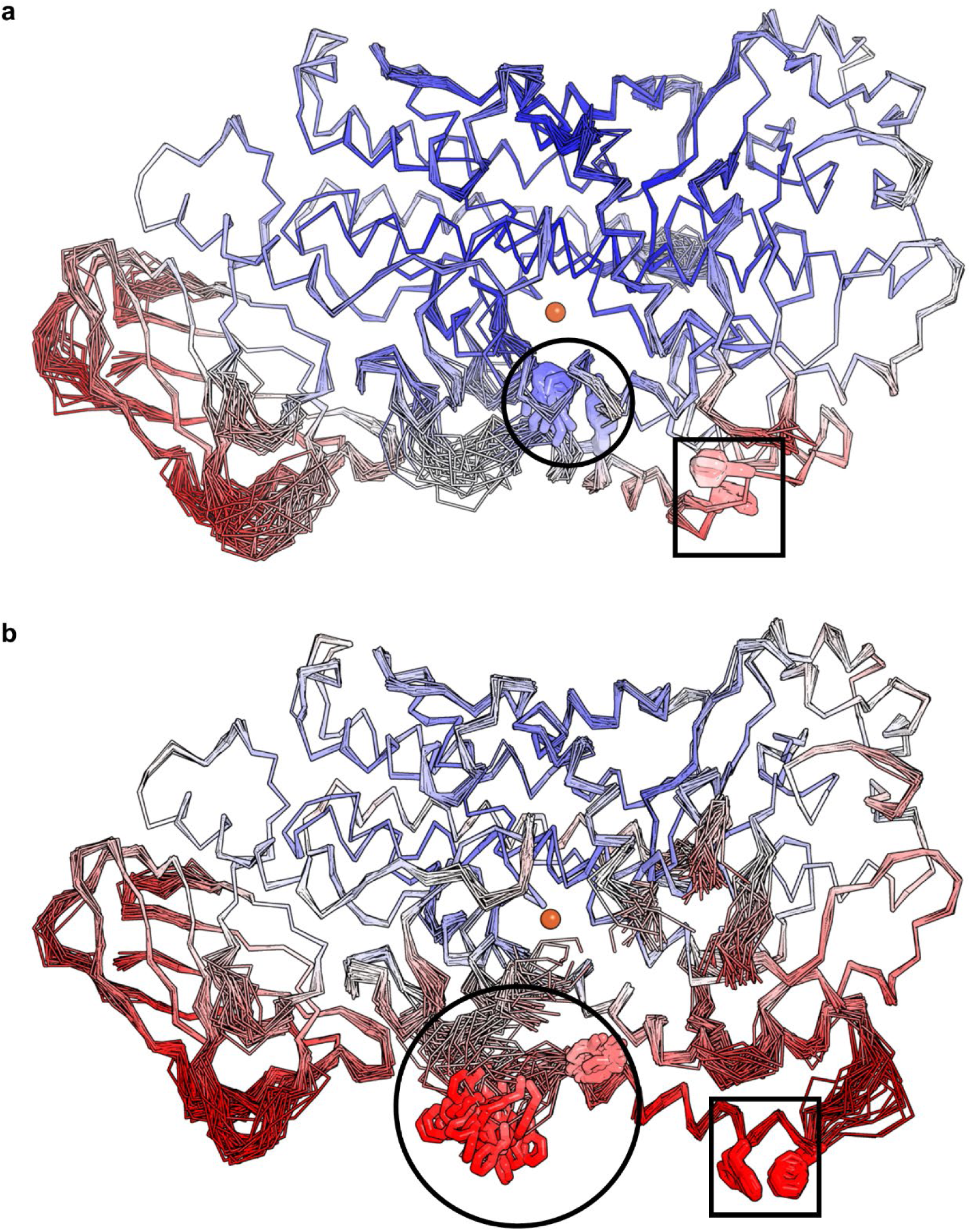
Ensemble refinement. **a**, Backbone trace of Stable-5-LOX in the closed conformation colored by B factors with blue as low, white as average, and red as high. The orange sphere denotes the catalytic iron. The average ensemble of the FY plug (blue, sticks in black circle) near the iron (orange, sphere) blocks the entrance of the substrate access portal. The backbone positions upstream (left) of the FY plug on Hα2 follow dissimilar paths consistent with flexibility. **b**, Stable-5-LOX in open conformation results in F177 of the FY plug (red, sticks in black circle) adopting multiple conformations around a flexible backbone. The two Phe residues (red, sticks in black box) at the C-terminus of Hα2 are relatively anchored as compared to the N-terminus of the helix.

### Mutants of Stable-5-LOX

As conformational flexibility may be a feature of 5-LOX, we next asked how these conformations of Hα2 affect enzymatic stability and activity *in vitro*. We prepared variants of Stable-5-LOX to favor the open and closed conformations by making substitutions that should (1) elongate Hα2 at the N-terminus, (2) “uncork” the active site, and (3) lock the C-terminal portion of the helix in “open” or “closed” conformations. G174 and D176 should favor the closed “broken” helix structure, as Gly is a helix breaker and D176 participates in a network of H-bonds and ionic interactions anchoring this end of the α-helix. Each were replaced by Ala, a residue that favors helical structures (G174A:D176A; “N-Elongated”) (Fig. 2). Replacement of Y181 to Ala should open access to the active site (Y181A; “Uncorked”) (Fig. 2) as shown by Mittal et al^27^. F193 and F197 are buried in the closed structure and participate in pi-stacking interactions with W201 and W599 and van der Waals contacts with R594 and R596. Their replacement with Asp could create an ionic lock with the nearby Arg residues, while substitution with Arg might drive them to their exposed conformation in the open structure by repulsion. Substitutions of the pair with Ala removes any favorable interaction they may make when buried inside the protein. These mutations are referred to as “C-Locked-Closed” (F193D:F197D), “C-Locked-Open” (F193R:F197R), and “C-Unlocked” (F193A:F197A) (Fig. 2). We combined the N-Elongated and Uncorked constructs in an elongated and open variant, named “N-Elongated-Uncorked” (Fig. 2). Lastly, we merged all the open-favoring mutations to form “Penta-5-LOX” (G174A:D176A:Y181A:F193R:F197R). These seven variants were assayed for proteolytic stability, kinetic activity, and product fidelity as described below.

### Evaluation of impact of mutations on overall fold

We previously reported that disorder near the amino-terminal region of Hα2 increases susceptibility of Stable-5-LOX to protease^22^, and we adapted this assay to establish how the point mutations in Hα2 might impact susceptibility. Overall, most open-model mutants tended to be more susceptible to cleavage proximal to Hα2, while C-Locked-Closed, Penta-5-LOX, and Stable-5-LOX were relatively protease resistant (Fig. 4a). Stable-5-LOX and C-Locked-Closed retained > 90% of the original band intensities in conditions in which C-Locked-Open retained only ∼30% of its full-length band (Fig. 4b). Interestingly, in Penta-5-LOX, wherein C-Locked-Open mutations are paired with the other elongated-helix-favoring mutations, resistance to proteolysis was restored to that of Stable-5-LOX (Fig. 4a). The intensity of the ∼50 kDa band that associates with loss of the N-terminal ∼150 amino acids was also monitored and showed reciprocal trends with increasing intensities as the 75 kDa band intensity decreased (Fig. 4b). The paired t-test was used to compare results to Stable-5-LOX.

**Fig. 4.**
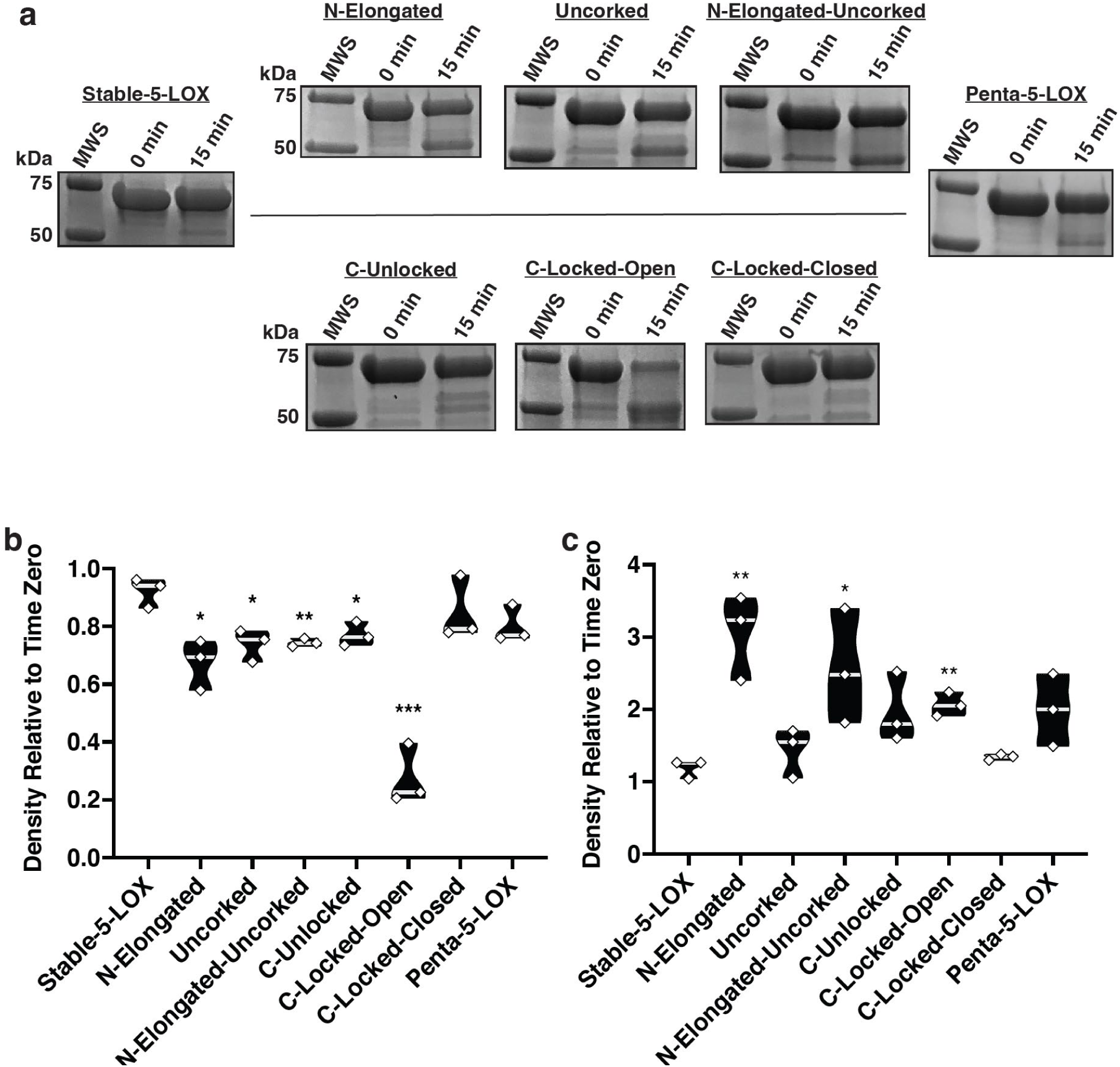
Evaluation of mutagenic impacts on overall fold. **a**, SDS-PAGE samples from each 5-LOX mutant. **b-c**, Triplicate measurements are shown in a violin plot with significant differences compared to Stable-5-LOX shown as astrics. *p<0.05, **p<0.01, ***p<0.001. Median is indicated by the light gray line. **b**, Densitometry of the full-length band after end-point normalized to time zero density. **c**, Densitometry of the ∼ 50 kDa band after end-point normalized to time zero density. MWS, molecular weight standard; 0 min, time 0 sample of pepsin digest reaction; 15 min, end-point of pepsin digest reaction.

Animal LOXs are named by their product regiospecificity with AA; hence 5-LOX generates primarily 5-hydroperoxyeicosatetraenoic acid (5-HPETE) and LTA_4_ to a lesser extent. We asked whether additional mutations along Hα2 would result in a major loss of product fidelity by the production of other HPETE isomers. HPLC analysis of the products formed by Stable-5-LOX and its variants indicate that the mutations did not display a major shift in product regio-specificity. The 5-HETE products recorded for each of the mutants meets or exceeds what is observed for the progenitor enzyme Stable-5-LOX (Supplemental Fig. 3). Collectively, the measured 5-HETE and LT values support the notion that Hα2 mutations can enhance enzymatic activity without increasing product promiscuity.

### Pre-steady state lag phase of Stable-5-LOX and Hα2 mutants

Similar to other iron-dependent LOXs, 5-LOX must be activated from its resting state^28^ by oxidation of Fe^2+^ to Fe^3+^. In a cellular context, the “peroxide tone” plays a role in this process^29^. *In vitro*, the enzymes can be activated by fatty acid hydroperoxides, and thus 5-LOX is routinely pre-incubated with an 18-carbon analog of the product, 13-hydroperoxy-9Z, 11E-octadienoic acid (13-Hpode)^28^. Without incubation with 13-Hpode, a distinct lag phase is observed until enough HPETE accumulates to fully activate the enzyme by oxidation so that it can achieve steady state. We reasoned that 13-Hpode likely accesses the catalytic iron *via* the putative substrate portal. In an effort to amplify any differences between our “open” and “closed” mutants due to active site access, we did not preincubate our enzyme with 13-Hpode. We reasoned that an “open” conformation may lead to a shortened lag phase and monitored the pre-steady state kinetics of the variants while noting the duration of the lag phase as indicated by the onset of steady-state activity. In addition, steady-state initial velocities were determined for each of the variants. The onset of steady-state activity across all substrate concentrations tested occurred at 30 to 50 seconds for Stable-5-LOX and 35 to 50 seconds for C-Locked-Closed (Fig. 5). By contrast, steady-state onset occurred for Penta-5-LOX at 1-3 seconds and for N-Elongated and N-Elongated-Uncorked at 2-7 seconds across all AA concentrations (Fig. 5). C-Locked-Open, being comparatively distal from the active site, exhibited steady-state onset at 7 to 15 seconds across substrate concentrations. C-Unlocked showed onset at 8 to 30 seconds and Uncorked at 12 to 18 seconds (Fig. 5). In contrast, pre-incubation of Stable-5-LOX with 13-Hpode virtually abolished the pronounced lag shortening it to 1 second (Supplemental Fig. 1a). The loss of the lag phase by pre-incubation with 13-Hpode and little-to-no lag by open mutants suggests that the lipid hydroperoxide might also “prime” 5-LOX, perhaps by shifting the ensembles to a more open conformation in which the substrate has ready access to the catalytic machinery.

**Fig. 5.**
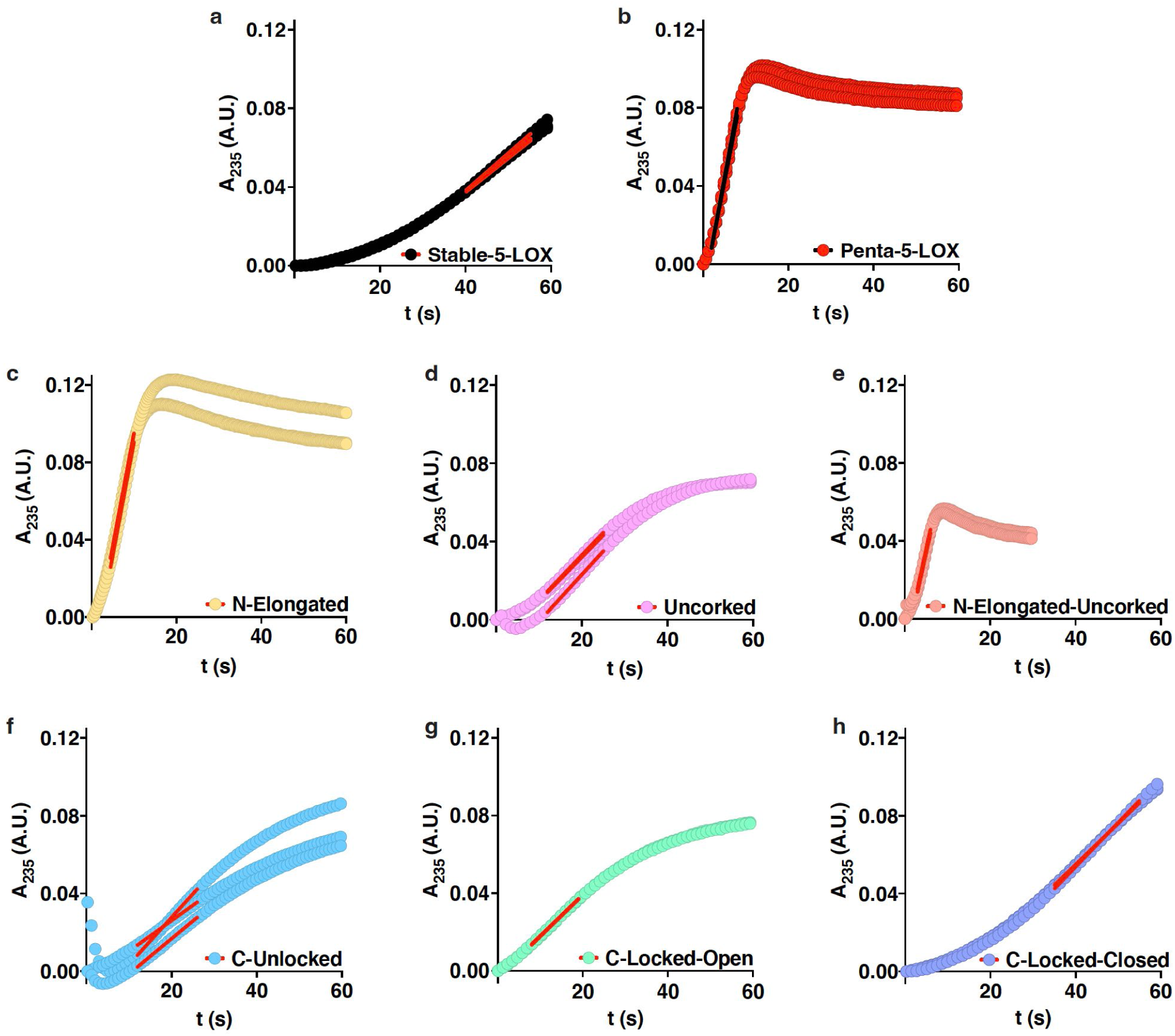
Comparison of kinetic lag phase among open and closed mutants. **a-h**, Absorbance at 235 nm vs. time plots; linear portions shown as red (black for **b**) lines. For all mutants, enzyme concentration was 250 nM and substrate concentration was 21.3 μM.

In summary, striking reductions in the lag time occurred when N-terminal Hα2 residues were mutated to favor an elongated helix and uncorking, while less dramatic reductions were evident for mutations introduced to favor C-terminal extension of Hα2. The data suggest that the N-terminal portion of Hα2 controls/impacts substrate access, but the conformational change that occurs to allow substrate entry likely extends beyond the putative substrate portal at Y181 to include distal amino acids at the C-terminus of Hα2.

### Kinetic parameters of Stable-5-LOX and Hα2 mutants

We found that plots of initial velocity vs. substrate concentrations are distinctly sigmoidal for Stable 5-LOX (Fig. 6a,b). A plot of AA concentration (μM) vs. initial velocity (v_0_, nM/s) exhibited a sigmoidal curve and is clearly a better fit to the Hill equation than the hyperbolic Michaelis Menton equation (Fig. 6a,b). Sigmoidal kinetics were previously reported for wild type human 5-LOX and attributed to interfacial interaction^30^. In a solution consisting of only phosphate buffered saline (pH 7.6), protein, and substrate, monomeric Stable-5-LOX yielded a hill coefficient of h = 2.11 ± 0.07 (Table 1). All Hα2 mutants fit the Hill equation and exhibited a sigmoidal curve with a hill coefficient greater than 1.7 (Table 1, Fig. 6a,b). Substrate affinity was maintained or slightly enhanced for Hα2 variants, showing a trend among mutants designed to favor an elongated Hα2 as having a higher affinity for AA (Table 1). Stable-5-LOX showed a k_cat_ of 0.43 ± 0.02 s^-1^. While k_cat_ was not significantly altered for C-Locked-Closed, C-Unlocked, C-Locked-Open, and Uncorked mutants; Penta-5-LOX, N-Elongated, and N-Elongated-Uncorked each exhibited a four-fold increase in k_cat_ (Table 1). This increase in catalytic activity by N-terminal Hα2 mutants in the absence of activating molecules provides insight into active site accessibility of Stable-5-LOX.

**Table 1.**
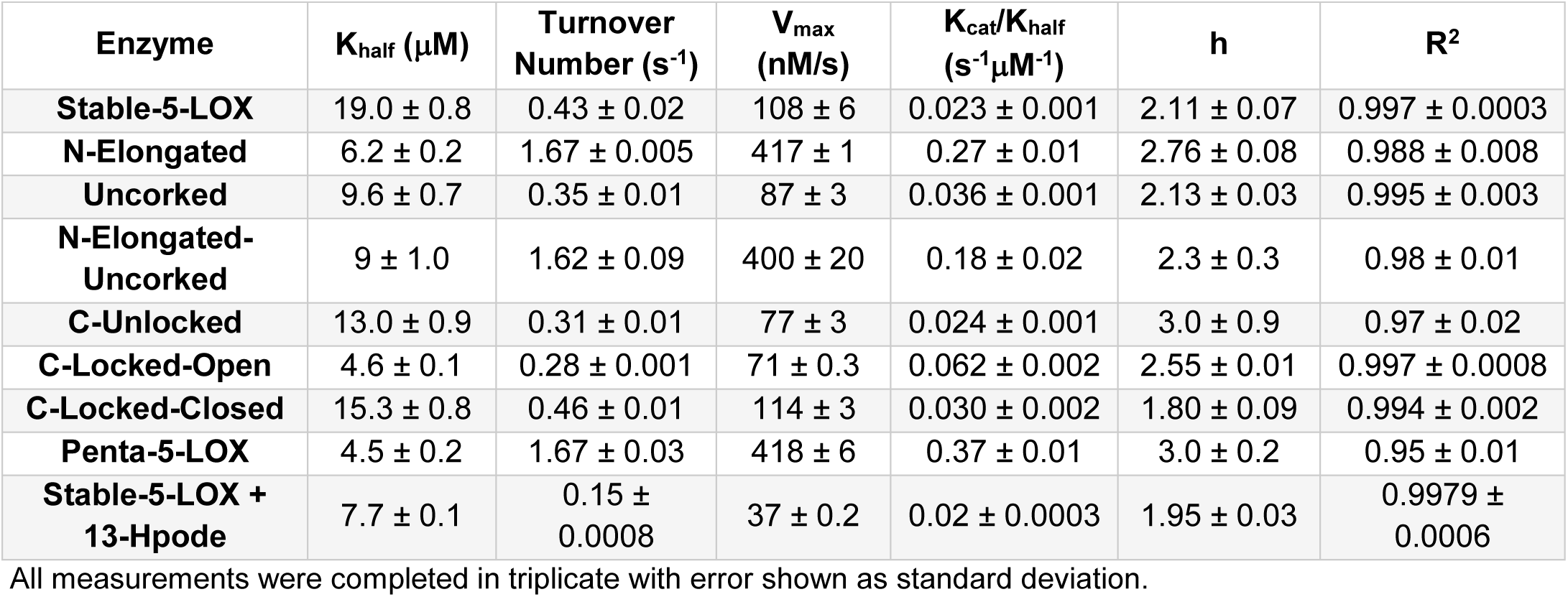
Kinetic parameters of open and closed mutants.

**Fig. 6.**
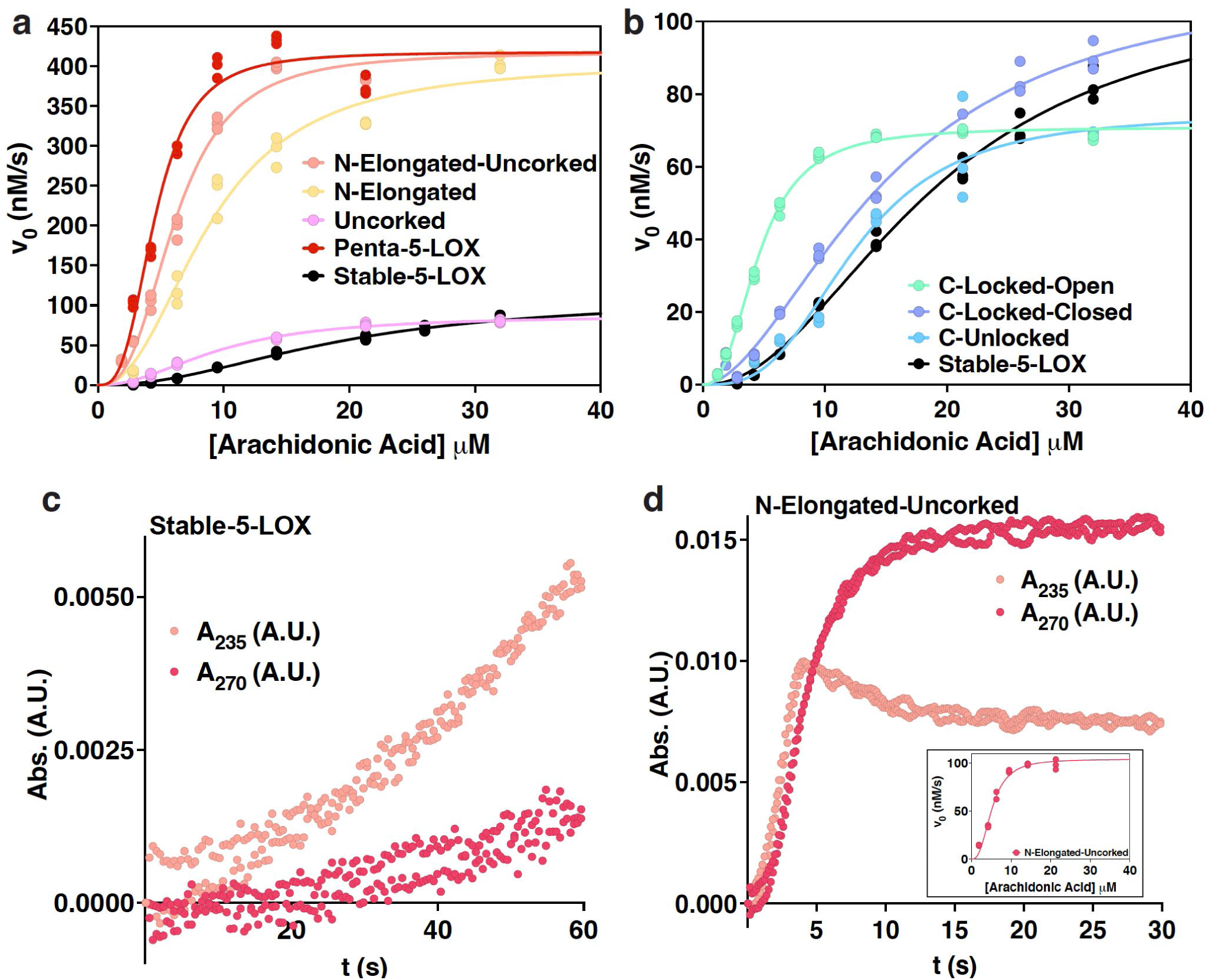
Kinetic characteristics of open and closed mutants. **a-b**, Saturation curves for Hα2 5-LOX mutants. Triplicate measurements were fit to the Hill equation. **a**, Amino-terminal Hα2 mutants compared to Stable-5-LOX. **b**, Carboxy-terminal Hα2 mutants compared to Stable-5-LOX. **c-d**, LT production by Stable-5-LOX and N-Elongated-Uncorked. Absorbance at 235 nm and 270 nm vs. time records HPETE and LT products, respectively. Duplicate measurements are shown for each with enzyme concentrations of 250 nM and substrate concentrations of 6.3μM. A saturation curve of LT production by N-Elongated-Uncorked produced by triplicate measurements is shown embedded in panel **d**.

When residues G174 and D176, which appear to resist helix formation, are replaced with Ala, the kinetic results indicate that substrate acquisition, product release, or both are enhanced. Note in Table 1 that k_cat_ was unchanged between N-Elongated and N-Elongated-Uncorked, but Fig. 5 shows that the duration of steady-state is shortened in N-Elongated-Uncorked compared to N-Elongated. Recall that 5-LOX catalyzes a second reaction—the transformation of the 5-HPETE intermediate to LTA_4_, which absorbs at 270 nm rather than 235 nm^31^. We asked if N-Elongated-Uncorked could also be rapidly performing the second reaction leading to an apparent shorter steady-state production of 5-HPETE. By monitoring intermediate and product turnover at both wavelengths, we observed an increase in absorbance at 270 nm that nearly parallels the increase at 235 nm in N-Elongated-Uncorked (Fig. 6d). This phenomenon is consistent with the idea that the 5-HPETE intermediate is being rapidly consumed for the production of the epoxide LTA_4_. The same robust activity was not found when observing Stable-5-LOX activity at the wavelengths mentioned (Fig. 6c). Interestingly, when Stable-5-LOX was incubated with 13-Hpode, the K_M_ was lowered to a similar range as Penta-5-LOX, N-Elongated, and N-Elongated-Uncorked (Table 1, supplemental Fig. 1). This observation is consistent with our hypothesis that 13-Hpode may shift the population of Hα2 conformers to the open conformation that AA can access with ease by out-competing 13-Hpode.

### AlphaFold2 prediction of Stable-5-LOX and variants

In the absence of structural models and despite extensive crystallization trials for the kinetically faster variants of Stable-5-LOX, we considered computational experiments to fill the gap. During the preparation of this manuscript, the structure prediction method AlphaFold2 (AF2) was published and showed predictive scores near atomic accuracy^32^. Subsequently, the ColabFold^33^ notebook was released, which allows novice users an opportunity to predict protein structures in the AF2 environment coupled through the Google Collaboratory cloud services. We utilized the advanced version of AF2 where no homology templates are allowed. While these programs are designed to predict overall structures rather than the impact of point mutations^34^, given the apparent structural heterogeneity of Hα2 in the context of a highly conserved fold, we asked whether the software might distinguish among the various substructures available to Hα2. We observed a trend in which those mutations that confer improved kinetic parameters and protease susceptibility are predicted by AF2 to statistically favor the open conformation to a greater extent.

We generated ten predicted models for each variant with all models scoring above 90 in the predicted local distance difference test (pLDDT). As expected, the structures of the membrane-binding loops on the β-barrel domain diverged more than the body of the protein (Fig. 7a) with some of the lowest pLDDT scores in these regions. Notably, structural divergence in the Hα2 region was consistently the highest with the lowest pLDDT scores. We superimposed all predicted models of the various mutants with the closed structure of Stable-5-LOX (PDB code: 3O8Y) to get an all atom r.m.s.d. given in angstrom (supplementary Table 2). We then visually inspected if Hα2 was in an elongated form and whether F177 and Y181 were plugging the active site closed (Fig. 7a, highlighted box). We classified the structures into three categories: closed (F177 superimposes with the 3O8Y.pdb), intermediate (FY plug has rotamer switched and moved 1-2 Å from the crystal structure), or open (FY plug is displaced similarly as in the open structure with an elongated conformation of Hα2). The progenitor Stable-5-LOX was predicted to have 8 closed, 0 intermediate, and 2 open conformers (Fig. 7b). In contrast, the mutant N-elongated is predicted to have only 1 closed, 2 intermediate, and 7 on a continuum of open conformers, consistent with a switch in the ensemble populations (Fig. 7c). Strikingly, the combination of uncorked and N-elongated is predicted to have all ten ensembles adopting a similar open conformation with little deviation of the backbone positions (Fig. 7d). This retrospective analysis of variants of Stable-5-LOX by AF2 agrees with our interpretation of the kinetic data: point mutations designed to favor the open conformation have higher rates of turnover. It will be interesting to see if other protein engineering efforts will capture sensitive point mutations responsible for conformational regulation by using the predictive power of the AF2 software.

**Fig. 7.**
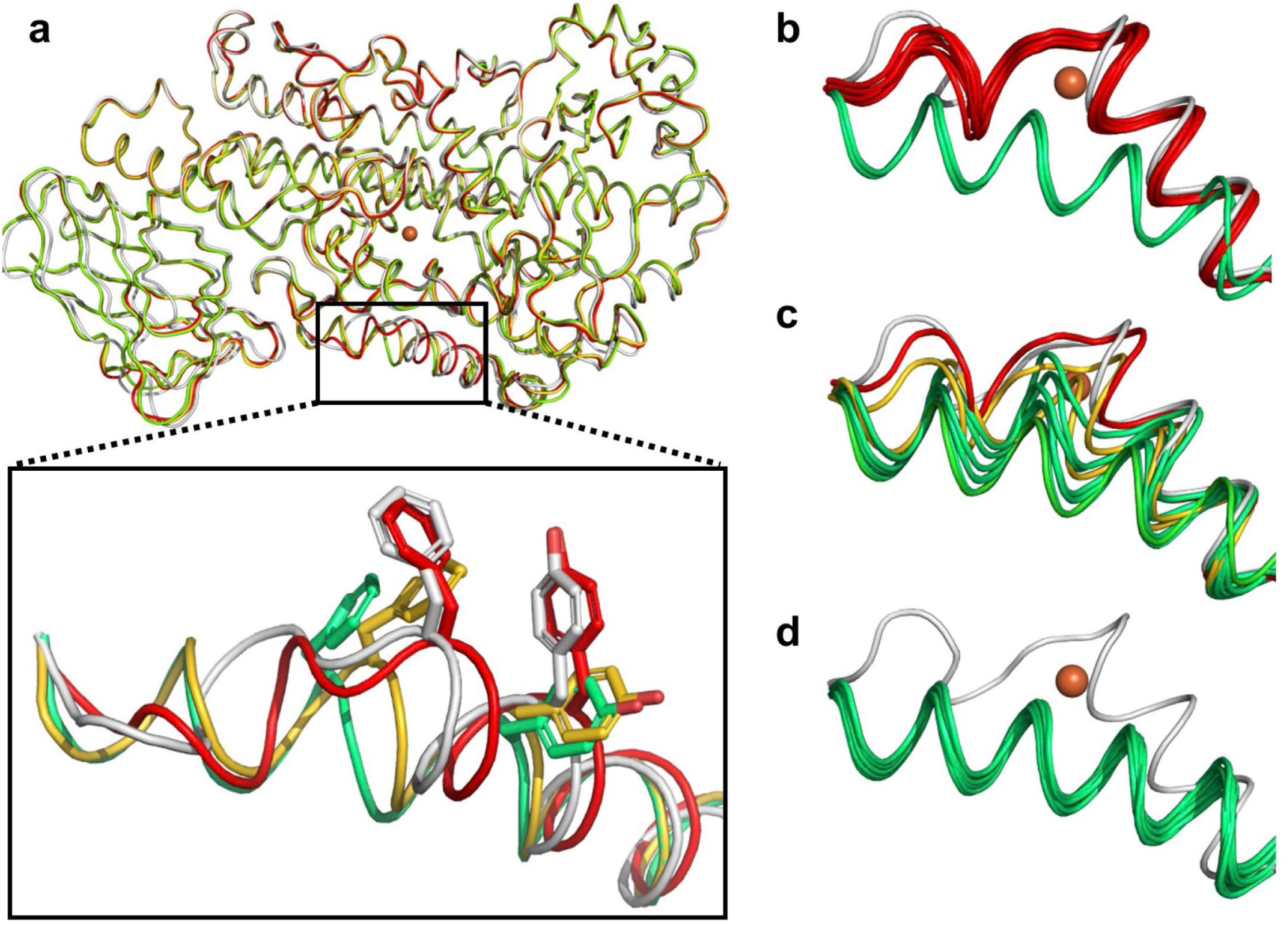
AlphaFold2-predicted structures of Stable-5-LOX and variants. **a**, Representative conformations (cartoon loop) of predictions from AF2 (closed conformation in red, partial open in gold, and fully open in green) superimposed on crystal structure (PDB code: 3O8Y in grey). Overall r.m.s.d. for predicted structures are ∼0.6 angstrom excluding regions of the β-barrel domain and more notably in Hα2 (highlighted in box below). The position of FY plug (side-chain, sticks) in respect to the substrate access portal was used for nomenclature of the conformation. **b**-**d**, Backbone of Hα2 of ten predicted structures for Stable-5-LOX, N-elongated, and N-elongated uncorked, respectively.

## Discussion

X-ray crystallographic studies revealed distinct conformational states of 5-LOX: one in which access to the catalytic machinery was blocked, and a second with unobstructed entry as a result of remodeling of a broken helical segment into a single, elongated helix. These two crystal structure “snapshots” in what is likely a dynamic ensemble continuum provide an excellent framework for understanding 5-LOX regulation through its interaction with substrate, membrane, and partner protein. The segmented Hα2 in “closed” 5-LOX remodels and elongates to adopt a structure similar to that observed for other “open” animal LOXs. The FY plug, which corks the substrate access portal, repositions to permit solvent access to the catalytic site. Ensemble refinement, where MD simulations are restrained by X-ray data, support conformational flexibility in these regions not only represented by high temperature factors but with multi-conformer ensembles. Experimental data, including proteolytic susceptibility and kinetic studies, support a model in which 5-LOX can adopt multiple conformations that differ in the structure of a helical segment that controls access to the active site. In lieu of crystal structures of the variants with these enhanced activities, predicted models by AF2 illustrate population-shifts in the ensembles consistent with our “open” and “closed” conformations. Although the AF2 algorithm has been considered unuseful for modeling the effect of missense mutations in other proteins (for a recent correspondence see^34^), these AF2-predictions of the various point mutants of 5-LOX with alternate populations of Hα2 positions may invigorate novel applications to other systems.

We were able to exploit the lag phase of Fe^2+^ activation in pre-steady state kinetic experiments. We correlated how different variants harboring mutations to favor the open conformation displayed shortened lag phases. Pre-incubation with the product analog 13-Hpode abolished the lag phase of the progenitor enzyme. However, because it likely requires the same access portal as substrate, we omitted this step to amplify any differences in access to the catalytic iron that may exist in the open vs. closed conformations. It is highly unlikely that lag phase variations are due to structural differences at the Fe^2+^-coordination sphere that might affect Fe reactivity, as the coordination spheres in the various animal and plant LOX structures^35^, whether open or closed or occupied with inhibitor or substrate or unoccupied, are virtually indistinguishable (for review see^18^). When open-conformation mutations were introduced along Hα2, significant alterations to turnover number, maximum velocity, and substrate affinity reflected a more active and easier-to-access enzyme machinery. This suggests that in a cellular context, in the absence of the Ca^2+^-signal that stimulates the release of AA from membrane phospholipids and the translocation of 5-LOX to the membrane, that the enzyme is mainly “closed,” with its active site protected. This hypothesis is supported by the earliest activation studies of 5-LOX in the 1980s that established that the addition of calcium^36^ and phospholipids^37^ in cell-free assays enhanced 5-HPETE and leukotriene production^38^, respectively. More recently, just the addition of exogenous AA,^39^ or more interestingly competitive inhibitors,^40^ is sufficient for the translocation of 5-LOX to the membrane, with the later hinting at membrane-binding determinants near the active site. We have potentially circumvented the requirement of these molecular activators by strategically removing the side chains required for restricting access to the catalytic machinery. Finally, human 5-LOX has been described as a labile enzyme with multiple modes of inactivation, including a turnover-based suicide inactivation^41^. A closed active site could provide protection from inactivation and limit the loss of the reaction intermediate before the enzyme can partner with FLAP and fully execute the two-step production of its potent inflammatory product. Moreover, in its open conformation the aromatic corking amino acids are poised for membrane interaction. Such an interaction may explain why substitution of F177 and Y181 with Ala has been shown to impair, but not abrogate, translocation of 5-LOX to the nuclear membrane in HEK cells^11^.

Kinetic data are consistent with AA shifting the population of Stable-5-LOX ensembles to the more active open conformation as revealed by sigmoidal curves of velocity vs. substrate concentration. Our model of a substrate-induced conformational change suggests that soluble Stable-5-LOX, a monomeric enzyme, displays cooperativity much like the wild type enzyme in the presence of an arachidonate/Tween 20 mixed micelle^30^. Cooperativity is generally associated with, but not limited to, oligomeric proteins (for review see^42^). Glucokinase, for example, is a monomeric enzyme that has one binding site for substrate but displays sigmoidal kinetics^43^. Cooperativity in monomeric enzymes may involve either an enzyme conformational equilibrium which is shifted in the presence of substrate, or a substrate-induced conformational change which slowly reassumes the inactivated conformation via hysteresis^44^ (for commentaries spanning from 1967 to 2015 see^45,46^). While our data cannot distinguish between these models, they demonstrate that neither the membrane nor FLAP are required to induce the change, yet clearly *in vivo* one or both factors may stabilize, or select for, the more active conformation. The lack of electron density for the arched helix along with the soaked substrate or inhibitors in the open 5-LOX structures could be due to the absence of the stabilizing partners of FLAP and the bilayer. An additional interpretation for the sigmoidicity observed may be that 5-LOX has an allosteric fatty-acid binding site, a mechanism proposed to explain the cooperativity observed for human 15-LOX-2^47^ and soybean LOX^48^. However, no unmodelled peaks in the electron density can be attributed to a hydrocarbon chain in the X-ray data sets. Apropos to a possible FLAP-stabilized interaction is the observation that one of the variants we studied (N-Elongated-Uncorked) could rapidly perform the second step and reach steady-state production of LT. This activity hints that an open conformation may be critical for the enzyme’s ability to rapidly convert 5-HPETE to LTA_4_ and leads us to speculate that its conformation, with Y181 displaced from the active site and the extension of Hα2, may mimic 5-LOX when interacting with membrane-embedded FLAP.

## Materials and Methods

### Protein expression and purification

The expression and purification protocol developed previously for Stable-5-LOX was utilized ^19^. Briefly, Rosetta 2 (DE3) cells harboring Stable-5-LOX in the pET14 plasmid were cultured in Terrific Broth and grown at 20 °C for 24 h. Cells were harvested at 6,000 g and frozen at -80 °C. Cell pellets were resuspended in BugBuster™ with protease inhibitors, and DNaseI was added before sonication and French Pressure cell lysis. TCEP (5 mM) was added to the clarified lysate to keep the protein reduced. Soluble material was applied to immobilized metal (Co^2+^) for affinity chromatography (HisTrap). The eluate was concentrated in 30 K cutoff (MilliporeSigma) and further purified by size exclusion chromatography with a Superdex-200 Increase 10/300 GL attached to an AKTA-FPLC (formerly GE LifeSciences now Cytiva). Protein purification was monitored by SDS-PAGE with ≥ 95% purity.

### Plasmid construction and mutagenesis

The Quikchange II XL Site-Directed Mutagenesis Kit (Agilent) was used to introduce point mutations contained in primer constructs. Mutations were introduced as follows: Uncorked, Y181A; N-Elongated, G174A + D176A; N-Elongated-Uncorked, G174A + D176A + Y181A; C-Unlocked, F193A + F197A; C-Locked-Closed, F193D + F197D; C-Locked-Open, F193R + F197R; and Penta-5-LOX, G174A + D176A + Y181A + F193R +F197A. All mutations were verified by sequencing.

### Crystallization

All structures reported are from soaking experiments of Stable-5-LOX crystals in the presence of substrate (AA) or inhibitors. More than 100 crystals were screened from soaking experiments for twelve different inhibitors or AA in aerobic or anaerobic environments, respectively. When AA was included in the experiments, the trials were performed in a Coy Anerobic chamber with the O_2_ level below 10 ppm. Thirty data sets were collected and eleven crystal structures refined with a remodeled helix α2. Protein crystals are grown by hanging-drop or sitting-drop vapor-diffusion at 295 K by mixing 1 μL of monomeric Stable-5-LOX (6-10 mg/mL) and 2 μL reservoir solution containing 7-12% Tacsimate, pH 6.0 (Hampton Research, Aliso Viejo, CA, USA). Crystals usually grow after 10 days and are subsequently transferred to a new sitting-drop vapor-diffusion plate with 50-70% Tacsimate (pH 6.0) and 0.2-1.0 mM inhibitor or substrate solubilized in DMSO for an 18 h incubation period. Crystals were vitrified in liquid nitrogen prior to shipment and data collection.

### Structure determination

Crystals were first screened at 100 K at the Protein Crystallography (PX) beamline at the Center for Advanced Microstructure and Devices (CAMD) at Louisiana State University. Full diffraction data sets were collected remotely at 100 K with a λ= 0.979 Å at the NE-CAT beamline 24-ID-C or 24-ID-E at the Advanced Photon Source (Argonne, IL, USA). Data were processed in the Rapid Automated Processing of Data (RAPD) software suite designed and supported by NE-CAT staff. Default strategies for data cutoff were allowed for data processing, with resolution being determined by CC_1/2_ cutoff of 0.35 or better for outer shell. Data were processed in space group P2_1_. For many datasets, we calculated a doubling of the unit cell volume (PDB code: 7TTL); a phenomenon reported in a previous structure of Stable-5-LOX (PDB code: 6NCF)^22^. After multiple rounds of refinement, we could clearly delineate a new conformation for Hα2 of Stable-5-LOX in one of the protomers of the asymmetric unit in both maps. However, electron density for the arched helix (414-429) and a neighboring loop (294-303) are disordered in this “open” protomer structure. Hα2 and a loop (170-216) becomes fully disordered in the other protomer of the asymmetric unit along with the arched helix (414-429), the neighboring loop (294-303), and the penultimate helix (594-604). A cutoff strategy for omitting peptides in areas of poor electron density was employed: (1) Is there no contiguous segment of density in the |2Fo-Fc| map observed at 1.0 sigma. (2) The B-factors in the missing peptide regions are ∼2X of the overall model. (3) The real-space correlation coefficients of the difficult peptide regions were < 0.5. Peptides that met these criteria were omitted from the model.

### Model building and phase refinement

Water, hydrogens, and the amino terminal histidine tag were stripped from a monomer of Stable-5-LOX (PDB code: 3O8Y) in Pymol 1.7^49^. Two or four monomers of Stable-5-LOX in the closed conformation were placed in the asymmetric unit by Phaser^50^ from the Phenix suite^51^ and the solution refined with Phenix.refine using rigid body, non-crystallographic symmetry restraints during early rounds of refinement, and real-space refinement^52^. A contiguous positive difference Fourier |Fo-Fc| in the Hα2 was observed in one of the protomers. A composite omit was generated with simulated annealing of the molecular replacement model and the newly observed conformation of Hα2 was modeled in the omit map. Real-space correlation coefficients for multiple peptide regions around the active site were below 0.5, and as described above omitted from the model.

### Ensemble refinement

The highest resolution structures of Stable-5-LOX in the “closed” conformation at 1.98 Å (PDB code: 7TTK) and the “open” conformation at 2.1 Å (PDB code: 7TTJ) were refined in phenix.ensemble.refinement^24^ in Phenix version 1.20.1-4487. The pTLS, which defines the fraction of atoms including in the TLS model, was setup using the different values of 1.0, 0.9, 0.8, and 0.6. We applied standard settings for all other parameters. The newest version of phenix.ensemble.refinement includes DEN restraints^53^, which allows for a more robust sampling of conformational landscapes^54,55^. The ECHT (Extensible Component Hierarchical TLS) disorder parameter was not installed for these current MD simulations restrained by X-ray data. The resulting ensembles for the closed structure includes 50 models (pTLS of 0.6) with the Rfree improving from 0.2011 to 0.1958. However, Ramachandran outliers increased significantly from 0.07% to a median of 3.02% and rotamer outliers increasing from 0.67% to a median of 12.52%. Most notable was the spurious placement of the catalytic iron in most of the 50 models. The open structure resulted in 40 models (pTLS of 0.6). The Ramachandran and rotamer outliers increased by 4.44% and 14.59%, respectively. Ensemble structures and maps can be found uploaded to Zenodo at https://doi.org/10.5281/zenodo.6037684.

### Proteolytic stability assays

Limited proteolysis assays were performed at 37 ºC for 15 minutes with a protein to pepsin ratio of 4:1^22^. Pepsin was prepared in NaAc (pH 4.0) at a concentration of 12.5 μM. Assays were performed in 0.1 M NaPO_4_ (pH 6.0) in the presence of ≤1.0% DMSO. The reaction was terminated by the addition of 2X SDS-PAGE loading dye mixed with 200 μM pepstatin A. Reaction samples were run at constant 150 V on Precast Mini-Protean 4-15% gradient TGX gels (BioRad).

Densitometry of the SDS-PAGE bands was performed using Adobe Photoshop 2020. Images were grayscaled and inverted in color, and levels were adjusted in order to maximize band clarity without removing pixels from the actual bands on the gels. Densities are reported as relative to that of the band at t=0. Background sections of exact selection sizes were taken for every measurement and subtracted from the densities. The paired t-test was used in GraphPad Prism to determine significant differences between Stable-5-LOX and the mutants.

### Fidelity assays

Product identification assays of AA oxygenated by 5-LOX and its mutants were performed in triplicate at room temperature. The total reaction volume was 1.0 mL with final concentrations of 200 nM Stable-5-LOX and/or mutant and 20 μM AA in 1X PBS (pH 7.6). Each reaction was run for 10 min and stopped with a volume of 650 μL of 100% MeOH and 30 μL of 1 M HCl. In addition, 10 ng of PGB_1_ was added to the sample to control for product extraction efficiency.

UCT Clean-UP^®^ columns were used for extraction under vacuum. Each sample was run on a separate column and washed with 1.0 mL of water followed by 1.0 mL of 25% methanol. Product was eluted with 300 μL of 100% MeOH, and the samples were dried under N_2_ and prepared for HPLC analysis by resuspension in 22 μL of 60% acetonitrile with 0.1% formic acid (pH 7.0). A small crystal of triphenylphosphine was added to each sample to reduce the remaining HPETEs to HETES and tubes were vortexed for 20 s. Under the standard enzyme assay conditions, HPETEs are rapidly reduced to hydroxyeicosatetraenoic acids (HETE). LTA_4_ is unstable; thus, its breakdown products are quantitated as four different leukotriene B_4_ isomers. Upon complete resuspension, the contents of each tube were injected into a Dionex ™ Multi-Wavelength High Performance Liquid Chromatography (HPLC) system using Thermo Dionex UltiMate^™^ ACC-3000 Autosampler for isocratic HPLC with 60% acetonitrile with 0.1% formic acid at 0.5 ml/min and elution was monitored with a diode array. HETEs are monitored at 235 nm and identified according to the elution times determined with standards. HETE isomers were quantified by area under the curve measurements divided by the PGB_1_ standard and molecular extinction coefficients. Simultaneous monitoring at 270 nm enabled measured of LTA_4_ hydrolysis breakdown products, identified by their characteristic trident structure at 270 nm.

### Kinetic assays

An Applied Photophysics SX20 stopped-flow spectrometer was utilized to monitor absorbance at 235 nm to determine kinetic constants for each of the variants. Triplicate measurements were taken for each concentration of AA, which ranged from 1.9 μM to 32 μM.

At least five concentrations of AA were used for each saturation curve. 1X PBS (pH 7.6) was used to perform assays with a final enzyme concentration of 250 nM. Faster mutants were tested at 100 nM final concentration as well to ensure no significant impacts of protein concentration on sigmoidal activity. AA was prepared by dilution to 48 mM in 100% ethanol and subsequent serial dilution in 1X PBS (pH 7.6) with ethanol concentration staying below 1% in the reaction mixture. Reaction times were between 30 to 120 seconds depending on the mutant. Stable-5-LOX was incubated with 2.5 μM 13(S)-Hpode for 10 minutes at room-temperature before triplicate measurements with AA for activator study.

GraphPad Prism was used to determine the slope, or initial velocity, of the most linear portion of A_235_ per second plots by simple linear regression. These velocities were then plotted vs. AA concentration and fit to various kinetic models. Data obtained at higher AA concentrations that are consistent with substrate inhibition, a phenomenon commonly observed in lipoxygenases, were not included in these fits. Based on R^2^, chi-squared values, and replicate tests for each data set, the allosteric sigmoidal fit was chosen for analyzing 5-LOX kinetics. R^2^ above 0.9400 and no significant deviation from the model based on the replicates test were used to evaluate the goodness of fit.

### AlphaFold2 predictions

We utilized the advanced notebook of ColabFold at https://colab.research.google.com/github/sokrypton/ColabFold/blob/main/beta/AlphaFold2_advanced.ipynb to make ten structural predictions of Stable-5-LOX and variants by the AlphaFold2. We also tested the original code from Deepmind, which resulted in structure predictions that differed little from those produced by ColabFold but required longer runtimes and produced fewer multiple sequence alignments (MSA). The MSA coverage was consistently above 3000 sequences for all positions when using the mmseqs2 method offered through ColabFold^56^. No homology templates were allowed and 5 models were predicted with either 3 or 24 recycles of structures fed back through the neural network for refinement. The pLDDT confidence values consistently scored above 91% for all models with little improvement in increased recycles from 3 to 24. The lowest predicted aligned error occurred in the membrane-binding loops and Hα2 (55-80%).

The low pLDDT in Hα2 was reflected in the heterogeneity of predicted structures in this region. All predicted models can be found uploaded to Zenodo at https://doi.org/10.5281/zenodo.6037684.

## Competing interests

All authors declare that they have no potential conflicts of interest.

## Notes

https://doi.org/10.5281/zenodo.6037684

### Competing Interest Statement

The authors have declared no competing interest.

