## Supplemental information and tables for "Helical remodeling augments 5-lipoxygenase activity"

**Supplementary Material**

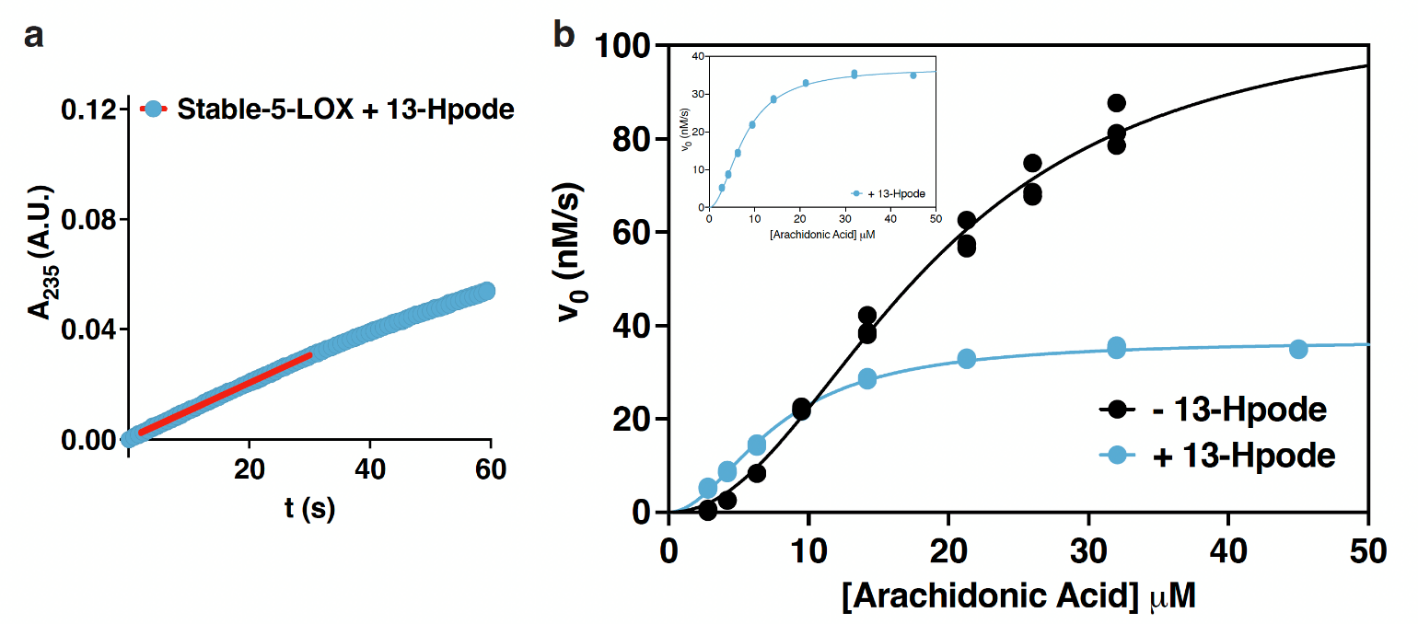

**Supplementary Fig. 1 | Stable-5-LOX activity with and without pre-incubation with the substrate analog 13-Hpode.** All enzyme concentrations were 250 nM and 13-Hpode concentration was 2.5 μM; each concentration of AA was tested in triplicate. **a**, Absorbance at 235 nm vs. time plot; linear portion shown as red. AA concertation was 21.3 μM. **b**, Saturation curves for Stable-5-LOX in the presence and absence of 13-Hpode. The embed­ded figure is given to show the shape of the curve in the presence of 13-Hpode.

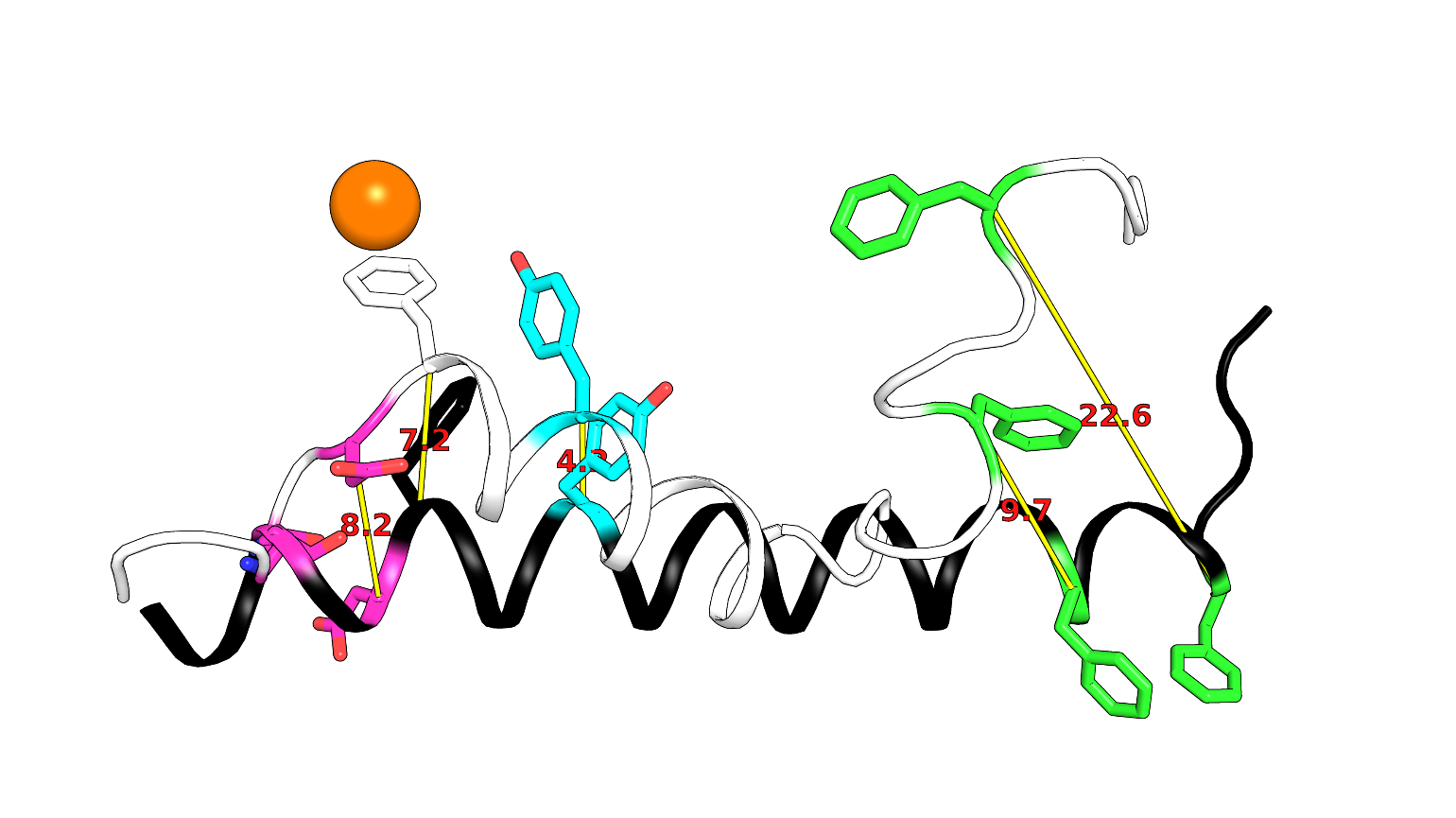

**Supplementary Fig. 2 | Displacement distances in Hα2.** Distances (yellow lines with red numbers between backbone positions of Elongated Hα2 (black, cartoon) and closed conformation (white, cartoon).

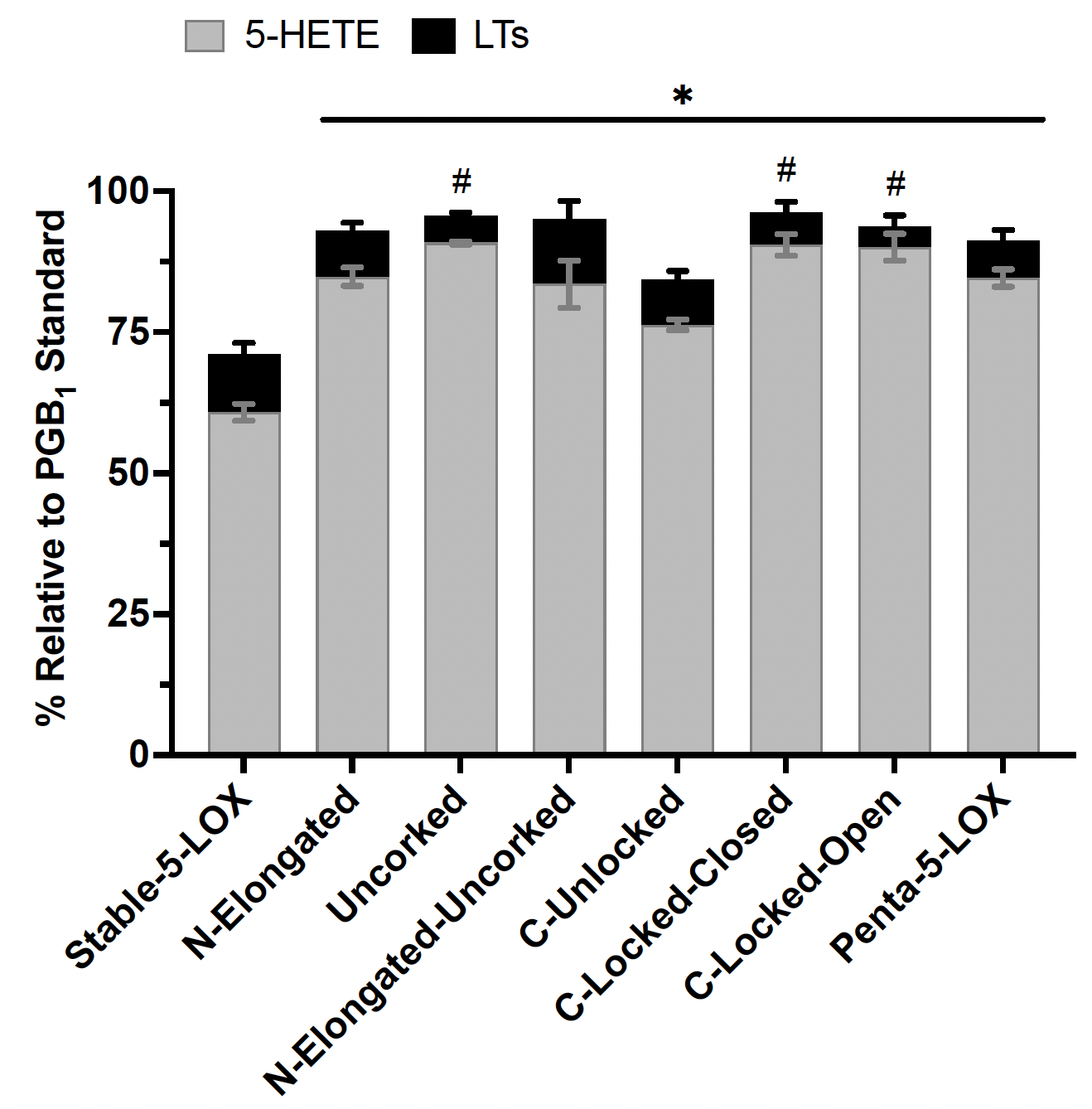

**Supplemental Fig. 3 | Verification of product fidelity among Hα2 mutants.** The formation of 5-HETE (grey) and LT (black) products was measured in triplicate for each mutant using Multi-Wave­length High-Performance Liquid Chromatography (HPLC). PGB_1_ was used as an internal stan­dard, and the percent of products were relative to the detected PGB_1_ concentration. PGB_1_ and LT concentrations were analyzed at 270 nm, while those for 5-HETE were evaluated at 235 nm. P-values are depicted as follows: 5-HETE, *p<0.05; LTs, #p<0.05.

### Supplemental Table 1.    Data collection and refinement statistics.

|  | Stable-5-LOX closed | Stable-5-LOX Open 2 copies | Stable-5-LOX Open 4 copies |
| --- | --- | --- | --- |
| PDB ID | 7TTK | 7TTJ | 7TTL |
| Wavelength | 0.9795 | 0.9792 | 0.9795 |
| Resolution range | 101.6 - 1.983 (2.053 - 1.983) | 50.25 - 2.1 (2.175 - 2.1) | 72.59 - 2.43 (2.517 - 2.43) |
| Space group | P 1 21 1 | P 1 21 1 | P 1 21 1 |
| Unit cell | 55.44 203.197 76.566 90 110.237 90 | 52.73 204.574 74.1155 90 107.65 90 | 76.791 204.547 106.768 90 109.035 90 |
| Total reflections | 502474 (42684) | 1116016 (58729) | 398092 (38837) |
| Unique reflections | 108039 (10149) | 86808 (8644) | 114424 (11353) |
| Multiplicity | 4.7 (4.2) | 12.9 (6.8) | 3.5 (3.4) |
| Completeness (%) | 98.43 (92.61) | 99.67 (99.12) | 97.91 (98.05) |
| Mean I/sigma(I) | 9.71 (1.20) | 18.81 (1.94) | 8.58 (1.16) |
| Wilson B-factor | 33.55 | 38.80 | 42.66 |
| R-merge | 0.1184 (1.177) | 0.1858 (1.007) | 0.1291 (1.064) |
| R-meas | 0.1335 (1.342) | 0.193 (1.091) | 0.1526 (1.263) |
| R-pim | 0.06044 (0.6301) | 0.05168 (0.4164) | 0.08058 (0.672) |
| CC1/2 | 0.996 (0.474) | 0.984 (0.802) | 0.996 (0.629) |
| CC* | 0.999 (0.802) | 0.996 (0.943) | 0.999 (0.879) |
| Reflections used in refinement | 107816 (10149) | 86542 (8600) | 114251 (11343) |
| Reflections used for R-free | 5264 (523) | 4058 (426) | 1975 (197) |
| R-work | 0.1652 (0.3173) | 0.1838 (0.3278) | 0.2156 (0.3532) |
| R-free | 0.2011 (0.3532) | 0.2225 (0.3598) | 0.2764 (0.3739) |
| CC(work) | 0.973 (0.784) | 0.960 (0.873) | 0.962 (0.772) |
| CC(free) | 0.962 (0.734) | 0.944 (0.794) | 0.921 (0.666) |
| Number of non-hydrogen atoms | 11684 | 10417 | 21687 |
| macromolecules | 10930 | 10056 | 21207 |
| ligands | 2 | 2 | 4 |
| solvent | 752 | 359 | 476 |
| Protein residues | 1345 | 1233 | 2607 |
| RMS(bonds) | 0.011 | 0.012 | 0.014 |
| RMS(angles) | 1.08 | 1.08 | 1.37 |
| Ramachandran favored (%) | 97.31 | 98.18 | 96.16 |
| Ramachandran allowed (%) | 2.61 | 1.82 | 3.84 |
| Ramachandran outliers (%) | 0.07 | 0.00 | 0.00 |
| Rotamer outliers (%) | 0.67 | 0.73 | 0.83 |
| Clashscore | 3.51 | 3.52 | 10.77 |
| Average B-factor | 43.17 | 56.84 | 60.09 |
| macromolecules | 43.15 | 57.14 | 60.38 |
| ligands | 39.26 | 47.87 | 41.73 |
| solvent | 43.52 | 48.37 | 47.63 |
| Number of TLS groups | 7 | 8 | 16 |

Statistics for the highest-resolution shell are shown in parentheses.

**Supplementary Table 2. AlphaFold2 statistics.**

| **Protein** | **RMSD** | **pLDDT** | **summary** |
| --- | --- | --- | --- |
| **Stable-5-LOX (3 cycles)** |  |  |  |
| **unrelaxed_1** | 1.299 | 94.64 | Closed |
| **unrelaxed_2** | 1.641 | 93.89 | Open |
| **unrelaxed_3** | 1.35 | 93.88 | Closed |
| **unrelaxed_4** | 1.31 | 93.25 | Closed |
| **unrelaxed_5** | 1.344 | 92.28 | Closed |
| **Stable-5-LOX (24 cycles)** |  |  |  |
| **unrelaxed_1** | 1.218 | 95.46 | Closed |
| **unrelaxed_2** | 1.304 | 94.68 | Closed |
| **unrelaxed_3** | 1.695 | 94.41 | Open |
| **unrelaxed_4** | 1.261 | 93.58 | Closed |
| **unrelaxed_5** | 1.277 | 93.17 | Closed |
| **N-elongated (3 cycles)** |  |  |  |
| **unrelaxed_1** | 1.558 | 94.31 | Open |
| **unrelaxed_2** | 1.549 | 93.87 | Open |
| **unrelaxed_3** | 1.593 | 93.83 | Open |
| **unrelaxed_4** | 1.462 | 92.41 | Open |
| **unrelaxed_5** | 1.493 | 91.22 | Partial |
| **N-elongated (24 cycles)** |  |  |  |
| **unrelaxed_1** | 1.529 | 94.38 | Open |
| **unrelaxed_2** | 1.332 | 94.15 | Closed |
| **unrelaxed_3** | 1.671 | 93.89 | Open |
| **unrelaxed_4** | 1.441 | 93.09 | Open |
| **unrelaxed_5** | 1.322 | 92.51 | Partial |
| **N-elongated-uncorked (3 cycles)** |  |  |  |
| **unrelaxed_1** | 1.679 | 94.16 | Open |
| **unrelaxed_2** | 1.643 | 94.12 | Open |
| **unrelaxed_3** | 1.665 | 94.06 | Open |
| **unrelaxed_4** | 1.528 | 92.66 | Open |
| **unrelaxed_5** | 1.633 | 91.46 | Open |
| **N-elongated-uncorked (24 cycles)** |  |  |  |
| **unrelaxed_1** | 1.63 | 94.79 | Open |
| **unrelaxed_2** | 1.607 | 94.53 | Open |
| **unrelaxed_3** | 1.59 | 94.35 | Open |
| **unrelaxed_4** | 1.564 | 92.78 | Open |
| **unrelaxed_5** | 1.629 | 92.19 | Open |
